# AMPGAN v2: Machine Learning Guided Design of Antimicrobial Peptides

**DOI:** 10.1101/2020.11.18.388843

**Authors:** Colin M. Van Oort, Jonathon B. Ferrell, Jacob M. Remington, Safwan Wshah, Jianing Li

## Abstract

Antibiotic resistance is a critical public health problem. Each year ~2.8 million resistant infections lead to more than 35,000 deaths in the U.S. alone. Antimicrobial peptides (AMPs) show promise in treating resistant infections. However, applications of known AMPs have encountered issues in development, production, and shelf-life. To drive the development of AMP-based treatments it is necessary to create design approaches with higher precision and selectivity towards resistant targets.

Previously we developed AMPGAN and obtained proof-of-concept evidence for the generative approach to design AMPs with experimental validation. Building on the success of AMPGAN, we present AMPGAN v2 a bidirectional conditional generative adversarial network (BiCGAN) based approach for rational AMP design. AMPGAN v2 uses generator-discriminator dynamics to learn data driven priors and controls generation using conditioning variables. The bidirectional component, implemented using a learned encoder to map data samples into the latent space of the generator, aids iterative manipulation of candidate peptides. These elements allow AMPGAN v2 to generate of candidates that are novel, diverse, and tailored for specific applications—making it an efficient AMP design tool.

## Introduction

AMPs contribute to the natural immune response in all classes of life and are active against a broad spectrum of microbes.^1,2^ Some AMPs are less likely to induce bacterial resistance, relative to traditional small molecule antibiotics.^3,4^ Additionally, AMPs can have synergistic effects when used in combination with traditional antibiotics^5–7^ or other AMPs.^8,9^

Over 15,000 antimicrobial peptides (AMPs) have been identified,^10^ but few have been advanced to clinical trials despite their promise as treatments for antibiotic resistant pathogens. Many known AMPs have limitations that have prevented effective therapeutic application, such as relatively low half-lives,^11,12^ undesirable or unknown toxicity to human cells, ^13,14^ and high production costs relative to traditional antibiotics.^13,15,16^

Designing AMP candidates that mitigate these shortcomings is a difficult problem. AMPs are made of amino acids arranged in a chain of arbitrary length, and feature a massive chemical search space. There are approximately 4.5 × 10^41^ unique peptides with 32 or fewer residues, if we consider only the 20 standard proteinogenic amino acids. Since the number of confirmed AMPs is low in comparison, it seems that the density of AMPs in the space of all peptides is also low. ^17^ Efficient methods are required to effectively develop AMP-based therapeutics.

Machine learning has aided in the discovery and development of AMPs, with many recent approaches relying on predictive models.^18–27^ Such approaches are usually labelled as quantitative structure-activity relationship (QSAR) models. The basic QSAR recipe is to select a property of interest (e.g., antimicrobial activity), train a machine learning model to predict that property using relatively easily obtained features (e.g., primary peptide structure), then apply the trained model to unlabelled samples to estimate the property of interest. After training, QSAR models can be used to identify properties of peptides present in a database that have yet to be experimentally validated.

The predictive approach can be extended to a generative one by adding an uninformed candidate generator (e.g., select a random peptide with length no more than 32). The randomly generated candidates can then be sorted and selected based on the property predicted by the QSAR model. This approach often suffers from excessive sampling requirements that inhibit discovery and design applications, due to the sparsity of AMPs in the peptide space. Additionally, reliance on engineered features constructed with domain expertise can further restrict the ability of these models to generate candidates that are qualitatively distinct from known AMPs. For example a commonly used feature is structure-based, however, at the time of writing only approximately 2.5% of known AMPs have structures, severely hindering the use of structure as an AMP predicting metric. In fact analyzing the presence of amino acids for structures with either alpha or beta characteristics (table S1 demonstrates that half the amino acids show up with less prevalance than chance, and of the ones that do show up a few show up in equal probability for each implying utility beyond structure. Even if the statistical rules were stronger, it is quite possible that some AMPs simply have no well defined structure.^28^

Explicitly generative models that are better informed by data can reduce the amount of sampling required to identify promising candidates. Proving this point, several studies have successfully applied recurrent neural networks (RNNs)^29^ ^30^ and variational autoencoders (VAEs)^31^ to AMP design and discovery.^32–36^ If we expand our scope to the more general case of molecular design, we find several more applications of VAEs,^37,38^ some of adversarial autoencoders (AAEs),^39,40^ and even the use of a generative adversarial network (GAN).^41^

Despite fairly broad adoption of machine learning techniques in this domain and growing interest in generative models, there is relatively little work investigating the use of generative adversarial networks (GANs) for AMP design and discovery.^42^ GANs are generative models that learn to produce samples from arbitrary data distributions by pitting a pair of artificial neural networks, dubbed the generator and discriminator, against each other in a zero-sum game.^43–45^ This family of models has seen great success in learning to generate images following an explosion of research interest in 2014.^45–50^ GANs can also generate text,^51–53^ a task that is qualitatively similar to AMP sequence generation and may indicate the potential for a new application.

Recently, we provided a proof-of-concept for such an application with AMPGAN and tested its ability to design antibacterial peptides. ^54^ For 12 generated peptides that are cationic and likely helical, we assessed the membrane binding propensity via extensive molecular simulations. The top six peptides were promoted for synthesis, chemical characterizations, and antibacterial assays. Three of the six candidate peptides were validated with broad-spectrum antibacterial activity.

GANs have served as core components in several creative image manipulation tools, ^55–57^ allowing for the generation of realistic looking images that satisfy user imposed constraints. Inspired by the iterative and controllable development process afforded by these creative image manipulation tools, we seek to apply similar models to AMP design. In particular, bidirectional conditional GANs (BiCGANs)^58,59^ are ideal for the AMP design task, since they provide a data driven generative process, designer control over some features of generated samples, and iterative development.

The data driven priors are learned via the zero-sum game between the generator and discriminator. In this game, the generator maps samples from a latent distribution (e.g., a multivariate normal distribution) to samples that appear to be drawn from the real data distribution, while the discriminator (or critic) is given samples and must identify if they were drawn from an authentic data distribution or produced by the generator. During training the discriminator minimizes a classification error, while the generator maximizes the error of the discriminator.

GANs can create realistic looking samples, but each sample will contain arbitrary features. In BiCGANs the control that we seek is created through the use of conditioning variables, ^46^ where the generator and discriminator are provided an additional input that contains metadata for the current sample. By allowing the discriminator to learn associations between features and conditioning variables, the generator is encouraged to account for the same associations, which then allows a designer to control the output of the generator. The conditioning variables are often constructed as binary vectors that indicate the presence or absence of the features of interest. For example, in an image generation context, a conditioning vector could indicate whether the generated image should contain certain objects.

The iterative development process that we want to enable is made possible by the bidirectional component of the BiCGAN. The bidirectional component is driven by a third network, the encoder, which maps data samples (e.g., AMP sequences) into the latent space of the generator. This allows real data samples to be projected into the latent space, which can be used to create landmarks in the latent space, facilitate latent space interpolations, and incrementally manipulate a particular sample.

In the following sections we discuss our training data, data pre-processing, and details of AMPGAN v2—our BiCGAN-based model for AMP design. We show that AMPGAN v2 can generate novel AMP candidates with similar physio-chemical properties to the training data, while also incorporating designer constraints.

## Methods and Models

### Training Data

We constructed our training set by combining the Database of Antimicrobial Activity and Structure of Peptides (DBAASP^10,60^), Antiviral Peptide database (AVPdb^61^), and UniProt^62^ databases. We extracted the FASTA formatted sequence information, microbe targets (e.g., Gram-positive bacteria, Gramnegative bacteria, viruses), mechanism targets (e.g., cell membrane, cytoplasmic protein, cell replication), and activity measures (primarily MIC50 measured in *μ*g/ml) from each database as available. Sequences containing non-FASTA symbols (e.g., tail modifications, lower case characters, etc.) or more than 32 amino acid residues were filtered. We chose MIC50 as our primary activity measure since it was one of the most prevalent measurements present in DBAASP. We did not consider other activity measures, such as MBC, due to difficulty in correctly combining such measurements with MIC50.

After removing duplicated sequences between DBAASP and AVPdb, as well as “false negative” sequences from UniProt that also appear in DBAASP or AVPdb, we obtained 6238 sequences from DBAASP, 312 sequences from AVPdb, and 490341 sequences from UniProt. If a particular sequence has measured effectiveness against multiple microbe targets or mechanism targets, then we considered the superset of these. For sequences that have multiple activity measurements against one or more microbes, all measurements with compatible units are converted to *μ*g/ml and the arithmetic mean was used to represent the general antimicrobial activity of the sequence.

#### Conditioning Data

We constructed conditioning vectors for our model using indicators for the target microbes, target mechanisms, MIC50 level, and sequence length (Figure 1). The target microbe classes are cancer, fungus, Gram-positive bacteria, Gram-negative bacteria, insect, mammalian, mollicute, nematode, parasite, protista, and virus. The target mechanisms are lipid bilayer, replication, virus entry, DNA/RNA, cytoplasmic protein, assembly, virucidal, membrane protein, surface glycoprotein, release, and unknown.

**Figure 1:**
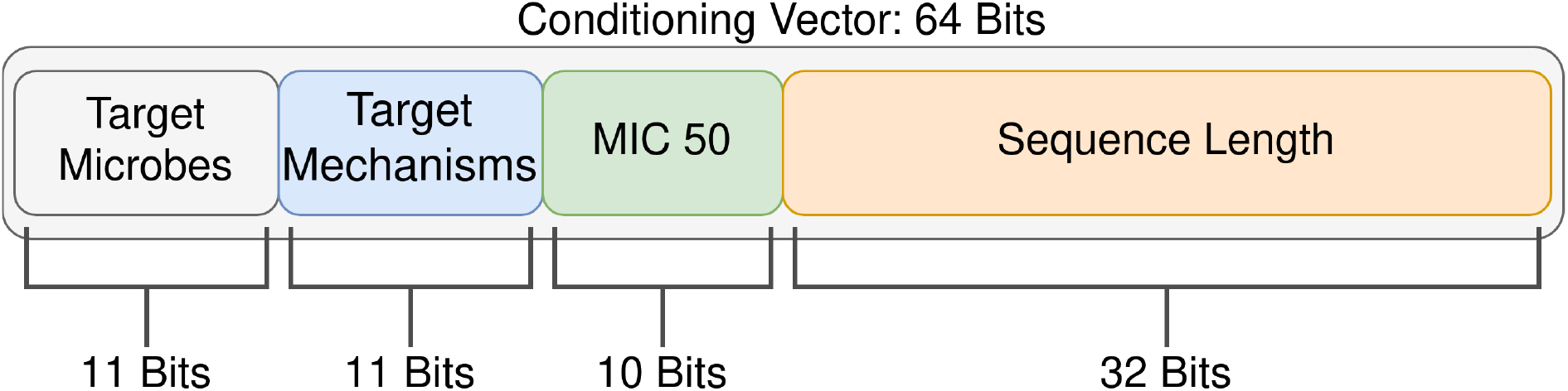
A visual summary of the contents and dimensions of a conditioning vector. All elements are binary encoded. For the target microbes and target mechanisms each element of the binary vector indicates activity against a particular microbe class or cellular mechanism. A one-hot encoding is used for the MIC 50 element, indicating membership in single MIC 50 decile. The sequence length is encoded as a bit mask, where 1 indicates the presence of a character and 0 indicates an empty slot.

The conditioning vector is then constructed as a 64 bit binary vector. The target microbes are encoded with 11 bits indicating activity, or lack thereof, against each microbe group. Likewise, the target mechanisms are encoded with 11 bits indicating interaction with a particular cell process or element. The MIC50 values are discretized into deciles using the following bin edges: 3.7 × 10^*−*6^, 5.7557 × 10^0^, 1.1 × 10^1^, 1.79869 × 10^1^, 2.7 × 10^1^, 3.88498 × 10^1^, 5.75996 × 10^1^, 8.53173 × 10^1^, 1.28 × 10^2^, 2.324687 × 10^2^, and 1.1240 × 10^4^ *μ*g/ml. Finally, the length of the sequence is represented using 32 digits, each indicating the presence or absence of a FASTA character.

We assumed that the sequences from UniProt did not have antimicrobial activity, since arbitrary peptides are unlikely to feature antimicrobial properties, and we already removed known AMPs. Thus, when we constructed conditioning vectors for these sequences the only non-zero elements were the length component, which was set appropriately, and the MIC50 component, which was set to the highest bin (lowest activity).

Figures S1 and S2 show the distributions of values across the conditioning vector elements (i.e. target microbes, target mechanisms, MIC50, and sequence length).

### AMPGAN v2 Design and Training

AMPGAN v2 is a BiCGAN constructed with three neural networks: the generator, discriminator, and encoder (Figure 2A).

**Figure 2:**
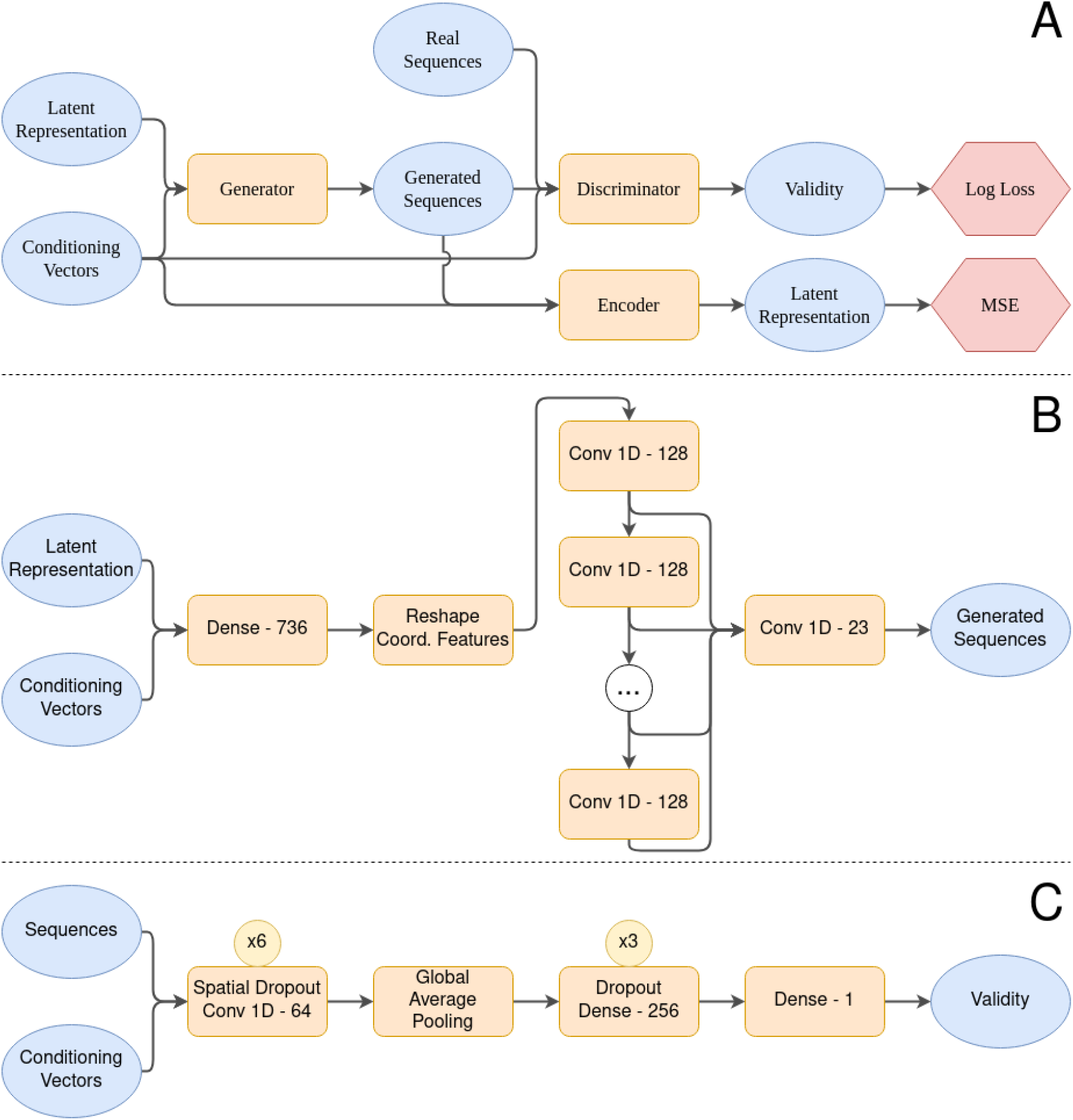
**A) AMPGAN v2 Macro-architecture.** AMPGAN v2 is a BiCGAN that consists of three networks: the generator, discriminator, and encoder. The discriminator predicts whether a sample is generated or not, and is updated using the log loss. The generator synthesizes samples, and is updated to maximize the loss of the discriminator. The encoder maps sequences into the latent space of the generator, and is trained using the mean squared error (MSE). **B) Generator architecture details.** We use 6 convolution layers in the central stack, each with a kernel size of 3 and an exponential dilation rate. All dense and convolution layers are followed by a leaky ReLU activation, except the final convolution layer, which has a hyperbolic tangent activation. The final convolution has a kernel size of 1. **C) Discriminator architecture details.** The convolutions use a filter size of 4 and a stride of 2. All applications of Dropout and Spatial Dropout use a drop rate of 25%. All dense and convolution layers are followed by a leaky ReLU activation, except the final dense layer, which has a sigmoid activation. The condition vectors are tiled and concatenated with the sequences along the features/channels dimension. The encoder uses the same architecture with a different output dimension on the final layer corresponding to the selected latent space dimension and a linear activation function.

The generator is composed of a dense layer that mixes the latent representation and conditioning vector, followed by a stack of exponentially dilated convolutions, and terminated by a single convolution that combines the multiscale features extracted by the prior convolution stack (Figure 2B). Global position information is added to the features as they enter the convolution stack to improve global sequence structure.^63^

The discriminator architecture contains a stack of strided convolutions, followed by several dense layers (Figure 2C). We apply spatial dropout before each convolution and dropout before each dense layer, excluding the output layer. Strided convolutions are used to quickly downsample the feature maps, while dropout increases the variance of the signal provided by the discriminator and can stabilize training. ^50^

The AMPGAN v2 encoder shares the same architecture as the discriminator, with the only difference being a larger final layer with a linear activation function.

We trained AMPGAN v2 for 2000 epochs, where AMPGAN v2 was shown all 6550 AMP sequences along with a random sample of the 490341 Non-AMP sequences in each epoch. Training proceeded with a batch size of 128 samples, where half were drawn from the AMP set and half from the Non-AMP set. The training signal for the generator and discriminator is provided by the binary crossentropy loss, while the mean squared error is used for the encoder. The discriminator is regularized using a gradient penalty, which has been shown to improve training stability and generalization. ^64,65^ In this configuration it takes roughly 30 seconds per epoch, adding up to 16 GPU hours for 2000 epochs using a Nvidia Tesla V100.

AMPGAN v2 builds on our previous experience with AMPGAN v1,^54^ though there are several differences in the implementation and evaluation procedure that make direct comparison of the two difficult. Full implementation details for AMPGAN v2 can be found in our GitLab repository. ^66^

## Results and Discussion

### Training Stability

GANs can be difficult to train depending on properties of their architecture and training data. Poor training stability can involve generator mode collapse, ^67–69^ cyclic generator-discriminator dynamics,^64,65,67^ and vanishing gradients caused by discriminator failures. ^70,71^

To investigate the training stability of AMP-GAN v2 we trained 30 replicates from scratch using different random initializations. We used a heuristic criteria with two conditions to determine if a trial is successful. First, the model must generate sequences with a character-level entropy that falls between 2 and 4. This removes models that tend to generate sequences with unrealistically low or high FASTA character diversity. For reference, the average character-level entropy across our training AMPs, non-AMPs, and their combination was ~2.6, ~3.43, and ~3.42 respectively. Second, the model must generate sequences whose length closely matches the value dictated by the conditioning vector. We quantified this by computing the *R*^2^ score over batches of generated sequences, and consider values greater than 0.5 to be successful.

These conditions were selected after observing two common failure modes in the training of AMPGAN v1. The first type were models that correctly handled the dictated sequence length, but only generated sequences composed of one or two amino acids. This resulted in a low character-level entropy, usually close to zero, and these models were clearly ineffective for generating true AMP candidates. The second failure mode resulted in models that produced sequences with more realistic character-level entropy, but completely failed to respond to the dictated sequence length. By not correctly responding to the elements of the conditioning vector, this type of model no longer provides human domain experts with a reliable method for directing the generative process, thus losing one of the primary benefits of the BiCGAN architecture that we have chosen.

We observed three successful trials that led to models with realistic sequence entropy and high correlation between the dictated sequence length and the length of the generated sequence. The other 27 trials failed to produce acceptable models, resulting in a ~10% training success rate. Figure S3 summarizes the variance observed during this experiment across several training metrics.

Our training success criterion requires that a successful generator account for the sequence length provided in the conditioning vector, but there is room for variation between the requirement of *R*^2^ = 0.5 and the ideal value of *R*^2^ = 1.0. Despite the allowed variance, all three successful trials resulted in models with high *R*^2^ scores–specifically 0.9852, 0.9986, and 0.9975. Qualitatively, this means that almost all of the generated sequences have a sequence length that is within ±3 of the dictated sequence length, which is visualized in Figure S4.

The observed ~10% training success rate increases the amount of resources required to train new iterations of AMPGAN, relative to a more stable model. Based on the estimate provided in the Design and Training section it will take an average of 160 GPU hours, a little less than a week, to obtain a quality model. However, this can be naively parallelized to reduce the wall clock time to only the 16 hours that it takes to train a single model.

Though it is inconvenient, the low training stability is not a dire issue, since an arbitrary number of AMP candidates can be generated once a quality model has been obtained. Also, It is likely that the training duration can be shortened from 2000 epochs to 1000 epochs, since Figure S3 indicates that all successful models had passed the criteria by that point.

We briefly investigated the training stability of our model on MNIST, an alternative dataset composed of handwritten digits. The digits were presented as a sequence of pixels, and the conditioning vectors were constructed using the classification labels. Under these conditions we found that our model trains quickly and reliably. This indicates that qualities of the training dataset may be the primary cause, rather than elements of the GAN architecture. We hypothesize that the lower quantity of labelled data and larger conditioning space of our training set (relative to MNIST) may contribute to the training instability.

### Physio-chemical Similarity

To be applicable to AMP design and discovery, we need to evaluate the quality of the generator and the properties of the generated candidates. However, it is prohibitively expensive to experimentally validate the ability of the generator to create sequences that follow the target microbe, target mechanism, and MIC50 values provided in the conditioning vector—so instead we focus on comparisons between easily measurable physio-chemical properties of generated and authentic peptide sequences.

We observe a high similarity between the amino acid distribution of the training and generated AMP sequences, which differ by less than 1% for most of the 20 natural amino acids (Figure S7). The most significant discrepancies come from Arginine (R) and Lysine (K), which are more prevalent in the generated sequences by 6.3% and 2.2% respectively. In contrast, three non-polar amino acids, Alanine (A) and Leucine (L) are 1.1% and 1.3% more common in the real AMP sequences respectively. Generally, these small differences suggest a consistency between the generated peptides and known AMPs. Figures S5 and S6 show additional amino acid distribution comparisons between various groups of peptides.

Figures 3 only investigates the appearance frequency of single amino acids, but there is a large body of research^72–76^ that suggests peptides feature complex grammatical structure. We investigated this higher-order organization using generalized word shifts,^77^ which extend the simple analysis done at the character level to sub-sequences of arbitrary length. Word shifts measure the contributions of distinct subsequences to a divergence measure between two groups of sequences and highlight the largest contributors.

**Figure 3:**
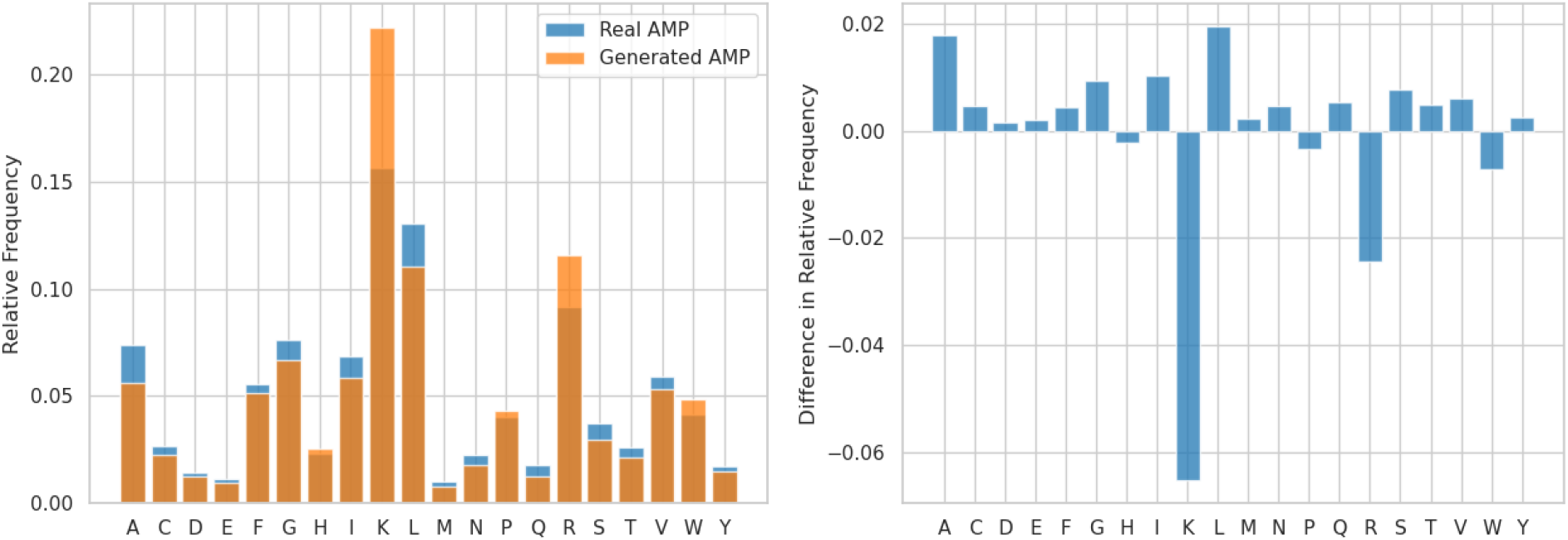
Distributions of amino acids present in generated vs non-generated AMP sequences. The distributions are layered in the left panel and the difference is shown in the right panel, facilitating different comparison perspectives. The generated distribution was created using 4855 sequences with conditioning vectors drawn at random from the training set. 50% of the conditioning vectors were taken from AMP sequences and 50% from non-AMP sequences. The model used to generate these sequences was arbitrarily selected from the set of successfully trained models. The nongenerated distribution was created using a sample of 5120 sequences that were randomly drawn from the training set with a 50%/50% split between AMP and non-AMP sequences. In all comparisons K is the largest outlier, appearing 4–6% more often in generated sequences than real sequences.

In Figure 4, we provide word shifts between generated AMPs and real AMPs for subsequences of length 2 and 3. The sub-sequences that were more common in generated peptides mostly involve one or more instances of K or R. Likewise the sub-sequences that were more common in real peptides tended to involve A or L. These two observations reinforce the results of the character level analysis. Many of the sub-sequences present in both plots feature positive charge or are hydrophobic, which corresponds well with known properties of alpha-helical AMPs. In the length 2 sub-sequence shift, the GP and PG motifs are of particular interest since they are often part of hinge-like structures near bends or kinks in proteins. Figures S7 and S8 provide baseline analysis that compares two uniformly randomly constructed samples of sequences using the same tools, which gives additional context for interpreting Figures 3 and 4 respectively.

**Figure 4:**
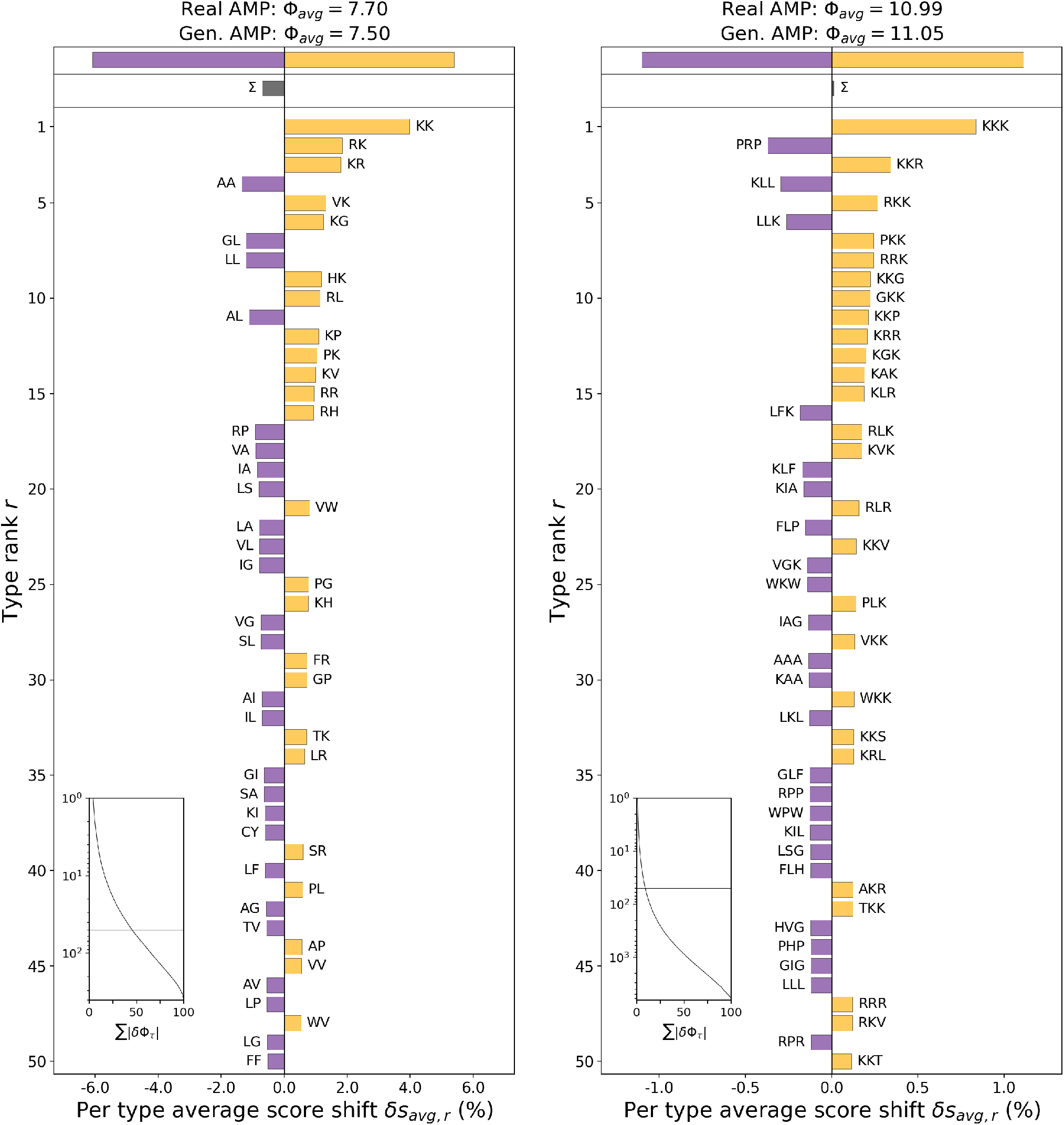
Shannon’s entropy divergence between the distributions of length 2 (left) and length 3 (right) sub-sequences of FASTA characters in AMPs from the training set (real) or AMPs created by the generator (generated). Purple bars indicate a greater prevalence of a particular sub-sequence in real AMPs, while gold bars indicate a greater prevalence in generated AMPs. The two values in the title of each panel indicate the average entropy of each group. For reference, the distribution of sub-sequences drawn from uniformly random sequences results in a maximum entropy of ~8.64 for length 2 sub-sequences and ~12.97 for length 3 sub-sequences. Both groups in both plots feature a lower entropy than the maximum, thus we should expect to see meaningful structures in each group. The CDF plot in the lower left corner of each panel indicates that the top 50 contributors to the divergence only account for ~50% (left) and ~10% (right) of the total divergence, thus both distributions are extremely flat.

### Sequence Diversity

When proposing candidate AMPs it is important that the generated candidates are diverse as a population and novel relative to known AMPs. If the generator produces sequences with low diversity, it can run into the same sampling problems as the extended predictive models discussed earlier. A generative model will be less useful for discovering new AMPs if it does not produce sequences that are novel relative to known AMPs. We applied the Gotoh global alignment algorithm^78,79^ to quantify the relative similarity of two bags of sequences. The distribution of alignment scores obtained between a pair of bags indicates the relative similarity of the bags, with more similar bags receiving higher scores.

Figure 5 contains letter-value plots^80^ that summarize the scores obtained by comparing the training AMPs, generated sequences, generated AMPs, and generated non-AMPs to themselves (i.e. a measure of diversity). Additionally, the final letter-value plot shows the distribution of global scores obtained by comparing the generated and training AMP sequences.

**Figure 5:**
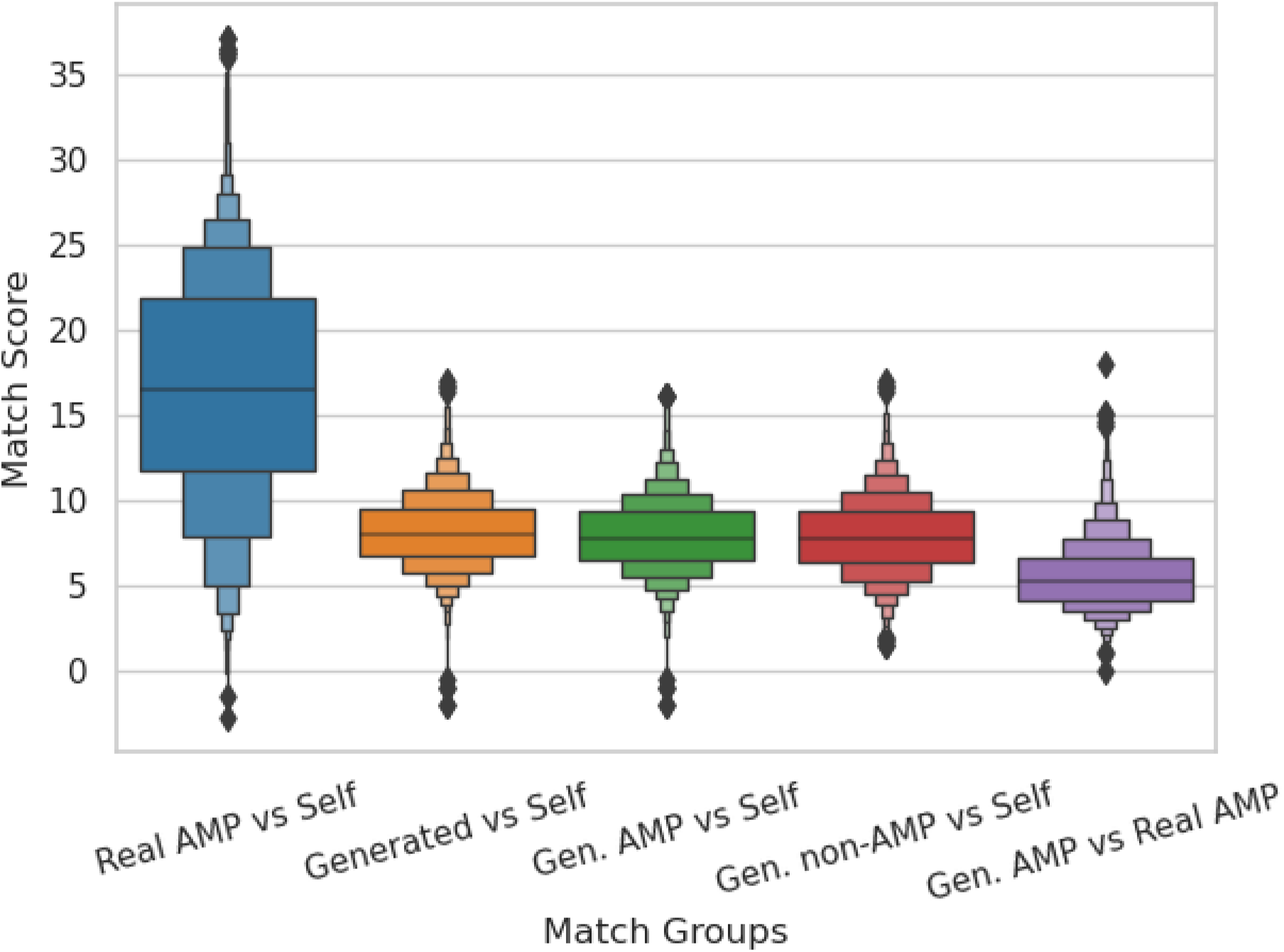
Letter-value plots showing distributions of match scores obtained from comparisons between different groups of sequences. The central horizontal line in each column denotes the median value. Each box extending from the median line indicates a percentile that is a half step between the starting percentile and the terminal percentile in that direction. For example, starting from the median line, the first box above is terminated at the 75th percentile, halfway between the 50th percentile and the 100th percentile. The diamonds in the tails indicate outliers, which in this case are approximately 5 to 8 of the most extreme values in each tail. The first distribution shows the match scores obtained when comparing the set of training AMPs with itself. The distribution of match scores for training AMPs has a median value that is approximately double that of the distribution for generated AMPs. This indicates that the set of generated AMPs is more diverse than the set of training AMPs. If we compare the generated AMPs directly with the training AMPs, which is shown in the final distribution, we find the lowest median match score observed so far. A low median match score here shows that the generated AMPs are novel relative to the training AMPs.

The training AMP score distribution features much higher median and upper percentile scores than any other distribution under consideration, indicating that there is relatively low sequence diversity in the training AMP set. The median score of 16.55 and mean score of 16.49 indicate a low diversity, especially relative to the generated AMP sequences that feature a median score of 7.83 and a mean score of 7.95. The generated non-AMP sequences feature a similar level of diversity to the AMP sequences, reaching a median score of 7.8 and a mean score of 7.92. The combined set of generated sequences obtains slightly higher scores than either the AMPs or non-AMPs separately, with a median of 8.0 and a mean of 8.17, which may indicate a slight chemical overlap between the two groups or may be due to chance. Comparing the generated AMPs with the training AMPs results in the lowest scores observed, with a median of 5.24 and a mean of 5.54, indicating that the generated AMPs are novel relative to the training AMPs. Figure S9 provides additional context for interpreting the global alignment scores shown in Figure 5.

### Estimated Antimicrobial Activity

We applied the predictive models developed by Waghu et al. to estimate the probability that sequences generated by AMPGAN will feature antimicrobial activity. This allows us to evaluate the quality of AMPGAN v2 in a absolute sense, ideally all AMP candidates generated by AMPGAN v2 would feature antimicrobial properties, and in a relative sense, by comparing it with AMPGAN v1.

We generated 5000 AMP candidates from AMPGAN v1 and 5000 from AMPGAN v2, then evaluated them using each of the four predictive machine learning models available on the CAMP_R3_ web page. From these predictions we calculated the percentage of sequences that were predicted to have antimicrobial properties, relative to the total number of sequences. Additionally, we estimated a 95% confidence interval for each percentage using bootstrapping. The results of this evaluation are summarized in Table 1, which shows that AMPGAN v2 strongly outperforms AMPGAN v1 which successfully predicted experimentally validated AMPs.

**Table 1:** Investigation of the expected antimicrobial properties of samples generated by AMPGAN v1 and v2 using the machine learning models developed by Waghu et al.. 5000 AMP candidates were drawn from each generative model and each candidate was evaluated by four predictive models: a support vector machine, a random forest, an artificial neural network, and discriminant analysis. The percentage of generated samples that were predicted to have antimicrobial activity is presented, along with a bootstrapped 95% confidence interval in parenthesis.

|  | AMPGAN v1 | AMPGAN v2 |
| --- | --- | --- |
| Support Vector Machine | 5.24% (4.44%, 6.08%) | 79.85% (78.27%, 81.39%) |
| Random Forest | 7.66% (6.68%, 8.72%) | 88.36% (87.06%, 89.62%) |
| Artificial Neural Network | 4.22% (3.52%, 5.00%) | 88.24% (86.94%, 89.46%) |
| Discriminant Analysis | 7.76% (6.76%, 8.72%) | 83.71% (82.23%, 85.18%) |

## Conclusion

In this work, we introduced AMPGAN v2, a BiCGAN that allows for the controlled generation of peptides with varying degrees of antimicrobial properties. We demonstrate that AMPGAN v2 can be trained successfully using a combination of AMP and non-AMP data. Notably, our data, from extensive comparison between known AMPs and generated peptides, indicates the capacity of AMPGAN v2 to generate sequences that are diverse and novel relative to the training data, but still maintain key AMP features. Additionally, AMPGAN v2 is responsive to changes in the conditioning vector, allowing for effective control of the generative process.

Based on the experimental validation of AMPGAN v1^54^ and the conditional VAE presented by Das et al., we expect the true success rate of AMPGAN v2 to be between 10% and 50%. If that proves to be the case, then AMPGAN v2 represents a fair improvement over the less than 1% success rate of more traditional design methods.^82^ Supporting this estimate, sequences generated by AMPGAN v2 were much more likely to be labeled as having antimicrobial properties than sequences generated by AMPGAN v1, when evaluated by a suite of predictive machine learning models.

AMPGAN v2 has many valuable features, though there are limitations that should be addressed in future work. Specifically, the low training stability of the current system should be improved to reduce training costs. Furthermore, additional validation is needed to ensure that AMPGAN v2 is responsive to manipulations of the target microbe and target mechanism conditioning elements. Greater responsiveness to manipulation of conditioning variables in combination with better training stability will improve designer confidence when developing new AMPs. Finally, additional quantitative methods for evaluating the quality of generative AMP models are needed to aid in development and performance comparisons. We believe that an extension of Fréchet Inception Distance^83^ to this domain and the use of Adversarial Accuracy ^84^ are promising directions to investigate. Along with these faster evaluation methods, we plan to experimentally validate the antimicrobial properties of several AMP-GAN v2 designed peptides.

AMPGAN v2 contributes a GAN-based model to an area where non-generative models or VAEs are more prevalent. Additionally, we open source AMPGAN v2, ^66^ allowing the community to interact with and deploy our tool to design and discover AMPs.

## Supporting Information Available

Distributions of conditioning variables, summary of training stability experiment, sequence length correlation figure, additional comparisons of amino acid frequency distributions, sequence analysis baselines, and global alignment score baseline.

## Data & Software Availability

All source code for the methods, experiments, and visualizations presented in this work are available under the MIT license via the project GitLab repository (https://gitlab.com/vail-uvm/amp-gan). All data required to train AMPGAN v2 is present in the GitLab repository, and can be obtained using the Git Large File Storage extension (https://git-lfs.github.com/).

## Acknowledgement

We thank Thayer Alshaabi, Lapo Frati, Gabriel Meyer-Lee, and Ollin Demian Langle Chimal for their helpful discussion and suggestions. Computations were performed on the Vermont Advanced Computing Core supported in part by NSF award No. OAC-1827314. JMR and JL were partially supported by an NIH R01 award (R01GM129431 to JL) and JBF was supported by an NSF award (CHE-1945394 to JL).

## Graphical TOC Entry

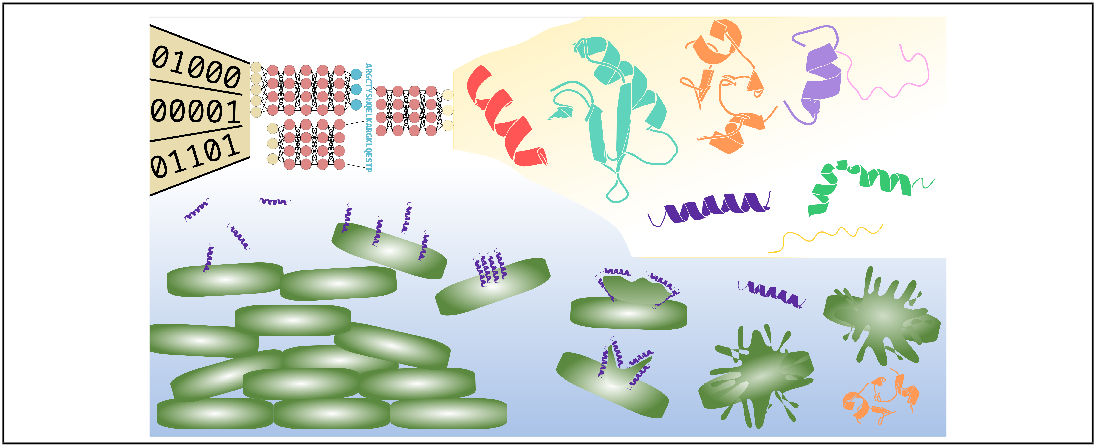

## Notes

https://gitlab.com/vail-uvm/amp-gan

https://dbaasp.org/

http://crdd.osdd.net/servers/avpdb/index.php

https://www.uniprot.org/

### Competing Interest Statement

The authors have declared no competing interest.

### Summary of Updates

Style and grammar polishing. Expanded discussion of the possible reasons for, impacts of, and solutions to the observed training instability. Added a new subsection to results, including a new table, which evaluated the expected antimicrobial activity of sequences generated by AMPGAN v1 and v2. Added SI availability statement and Data/Software availability statement.

