## Supplement for "AMPGAN v2: Machine Learning Guided Design of Antimicrobial Peptides"

### Supporting Information Available

#### Sequence Structure Profile

Table S1: Percentage of each amino acid's presence in the respective structure type. A completely random ordering should result in a table of 5% for all positions.

| Amino Acid | Helix | Sheet |
|---|---|---|
| A | 9.29 | 3.60 |
| R | 5.44 | 9.86 |
| N | 2.87 | 4.10 |
| D | 3.00 | 2.69 |
| C | 2.34 | 13.5 |
| E | 3.16 | 2.39 |
| Q | 2.65 | 2.59 |
| G | 10.0 | 10.8 |
| H | 2.38 | 1.54 |
| I | 7.13 | 4.59 |
| L | 10.9 | 4.69 |
| K | 12.8 | 5.87 |
| M | 1.42 | 0.88 |
| F | 4.98 | 3.70 |
| P | 3.14 | 4.16 |
| S | 5.41 | 6.42 |
| T | 3.39 | 5.41 |
| W | 1.87 | 2.00 |
| Y | 1.71 | 5.18 |
| V | 6.18 | 6.03 |

### Conditioning Information Distributions

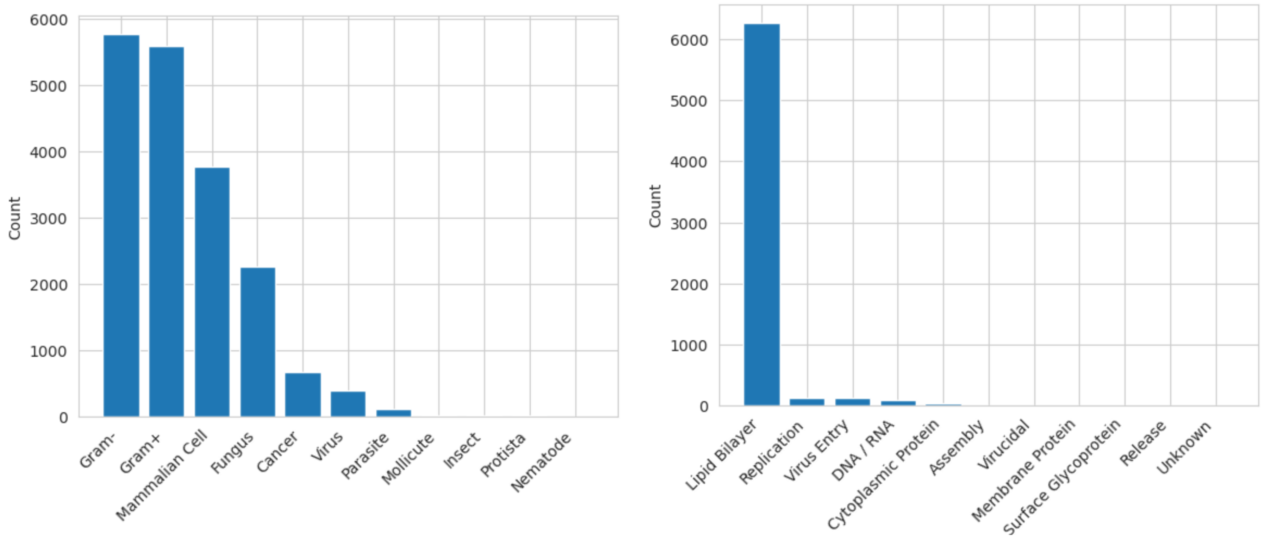


Figure S1: Label frequency for the target microbe (Left) and target mechanism (Right) conditioning variables.

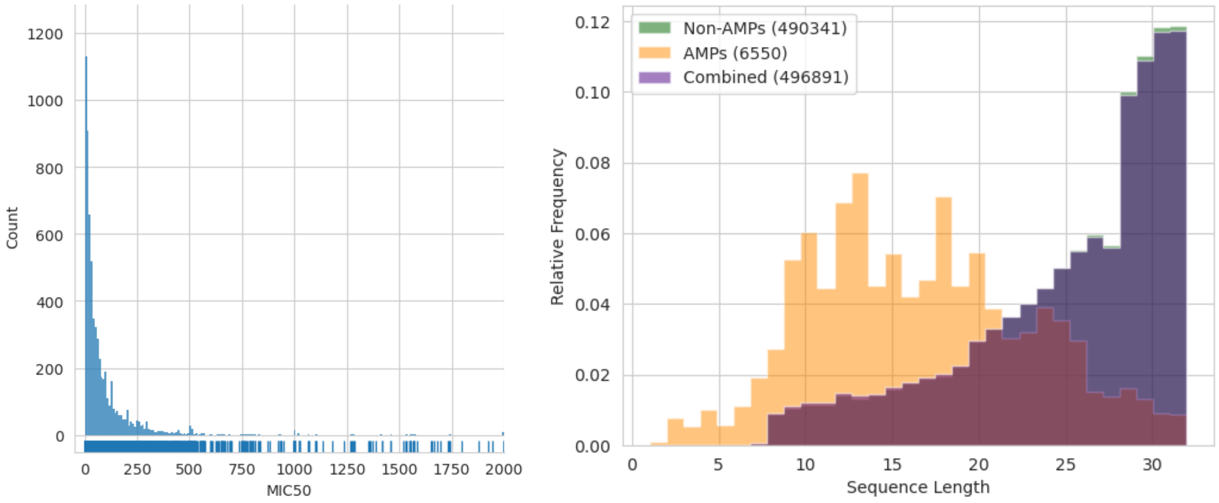


Figure S2: The distribution of MIC50 values before discretization (Left) and peptide sequence lengths (Right). 27 samples with MIC50 values greater than 2000 were truncated to ease inspection of the rest of the distribution.

### Training Stability

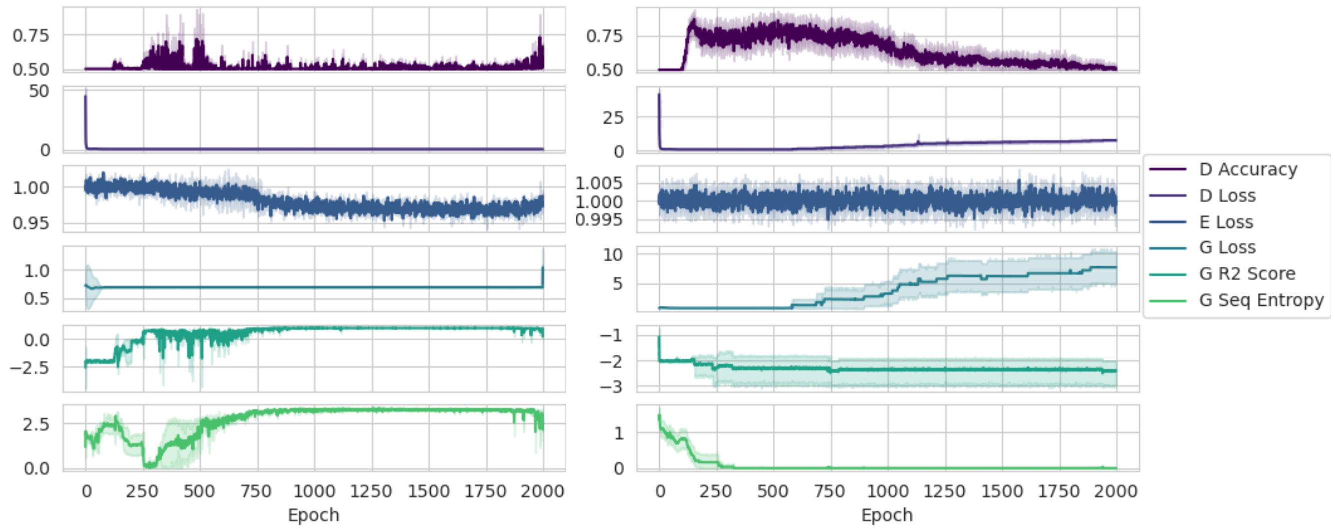


Figure S3: Investigation of training stability, summarizing the results of 30 independent trials. The left panel was constructed using the successful trials (3/30) and the right panel was constructed with the failed trials (27/30). From top to bottom the panels display the classification accuracy of the discriminator, the discriminator loss (log loss), the encoder loss (MSE), the generator loss (log loss), the $R^2$ score between the length dictated by the conditioning vector and generated sequences, and the average character-level entropy calculated over batches of generated sequences. This experiment highlights the relative instability of AMPGAN v2, with a success rate of $\sim$10%.

#### Sequence Length Correlation

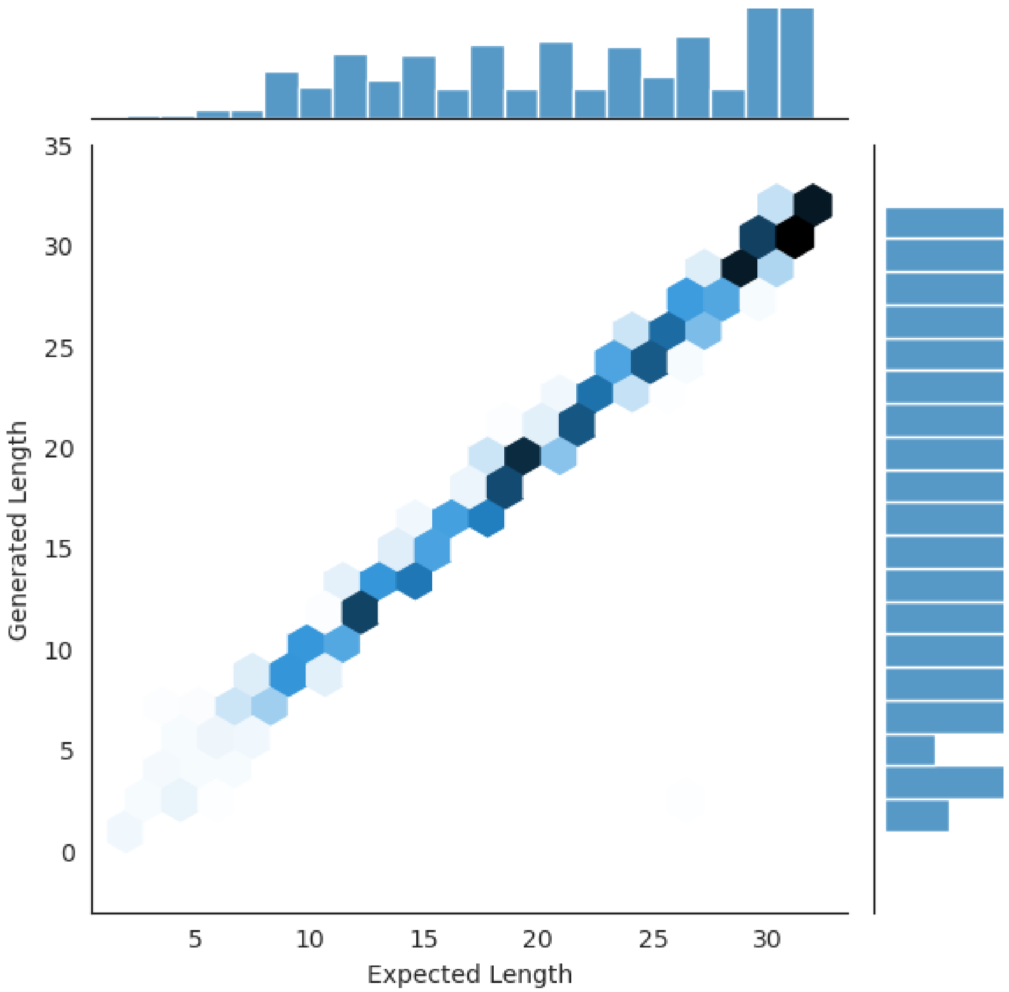


Figure S4: Agreement between the sequence length dictated by the conditioning vector and the length of sequences produced by the generator. This figure was created using 4855 sequences that were generated using conditioning vectors drawn at random from the training set. 50% of the conditioning vectors were taken from AMP sequences and 50% from non-AMP sequences. The model used to generate these sequences was arbitrarily selected from the set of successfully trained models. The generator pays close attention to the sequence length conditioning variable, resulting in an $R^2$ score of 0.9798.

#### Amino Acid Distribution Comparisons

Figures 3 and S5 only compare real and generated distributions, however, the relationship between AMPs and non-AMPs within each group is also important. The generator may create AMP sequences that are similar to real AMP sequences and non-AMP sequences that are similar to real non-AMP sequences, but fail to adequately capture the relationship between AMP and non-AMP sequences. To investigate this we create two additional comparisons between AMP and non-AMP sequences in both real and generated groups (Figure S6). Though there are some slight deviations present in the individual distributions in the panels on the left, which were already identified in Figure 3, the differences shown in the panels on the right are nearly identical. This indicates that the generator has learned the relative relationship between AMP and non-AMP sequences, despite some slight biases in its understanding of those distributions individually. Additionally, the lower

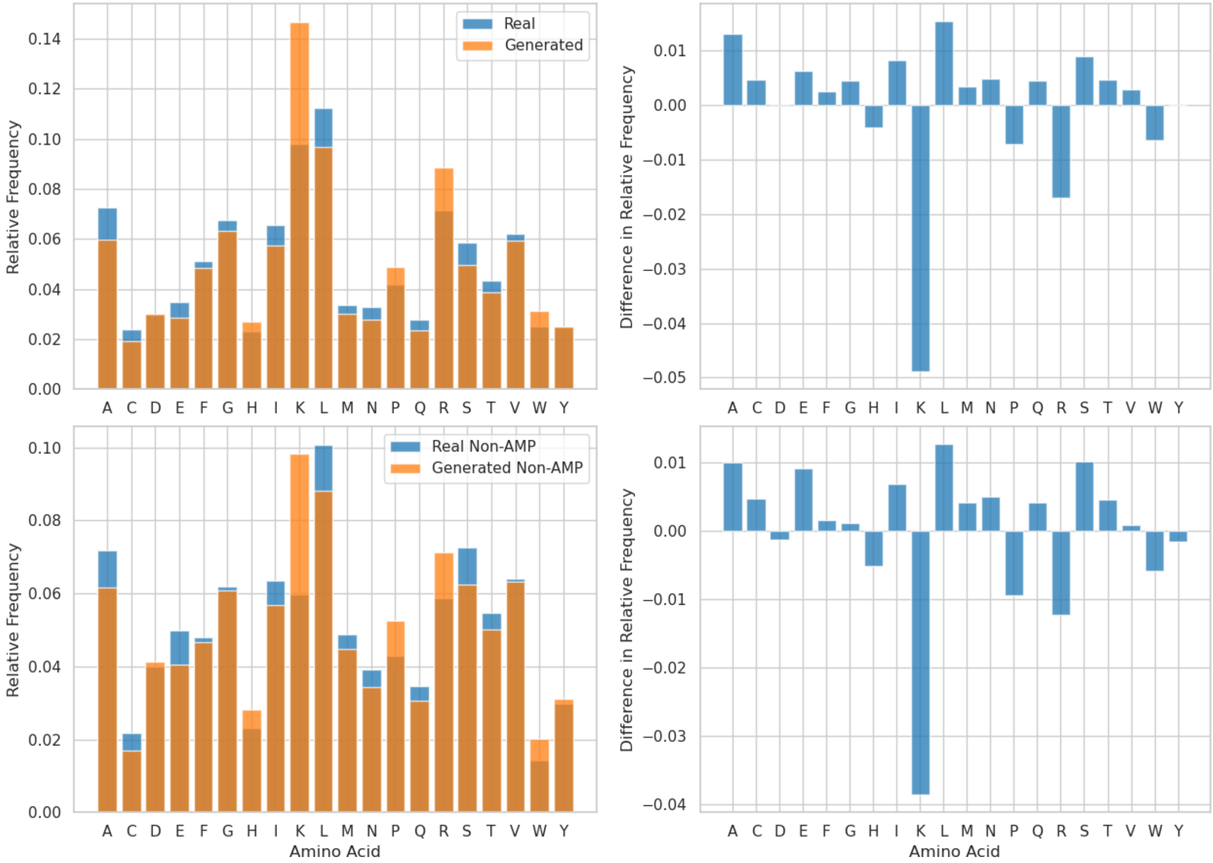


Figure S5: Distribution of amino acids used in generated vs non-generated sequences. Similar to Figure 3, but shows the distributions for all sequences (top) and non-AMP sequences (bottom). K remains the largest outlier, appearing 4-6% more often in generated sequences than real sequences.

right panel of Figure S6 is directly comparable to Figure 3 from Das et al., which agrees with the signs of the relative changes shown here for all amino acids except F and G. This qualitative agreement may indicate that both models have accurately captured the qualities of the training data distribution, or at least that both models acquired a similar bias profile.

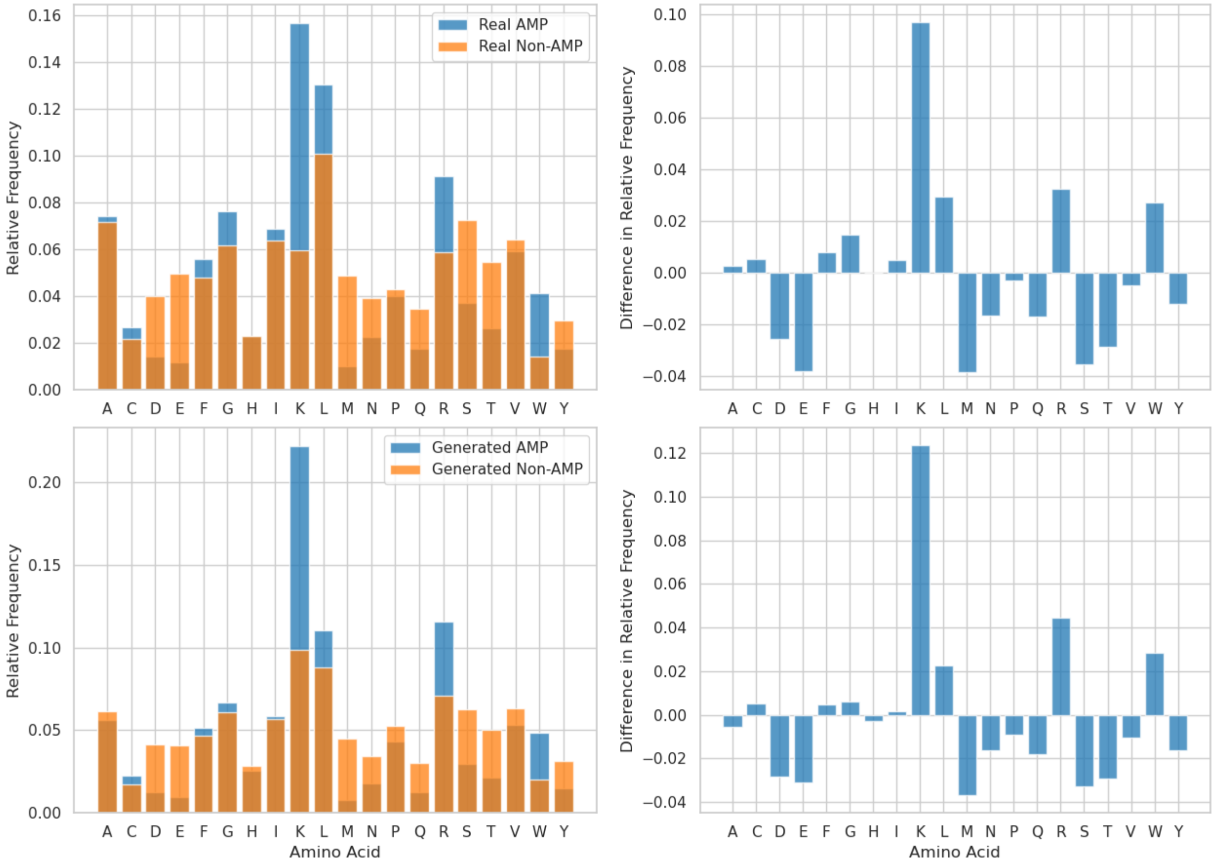


Figure S6: Amino acid usage frequency distributions for generated AMP and generated Non-AMP sequences (left) along with the difference between the two distributions (right). Comparisons are made between real (top) and generated (bottom) groups.

### Sequence Analysis Random Baselines

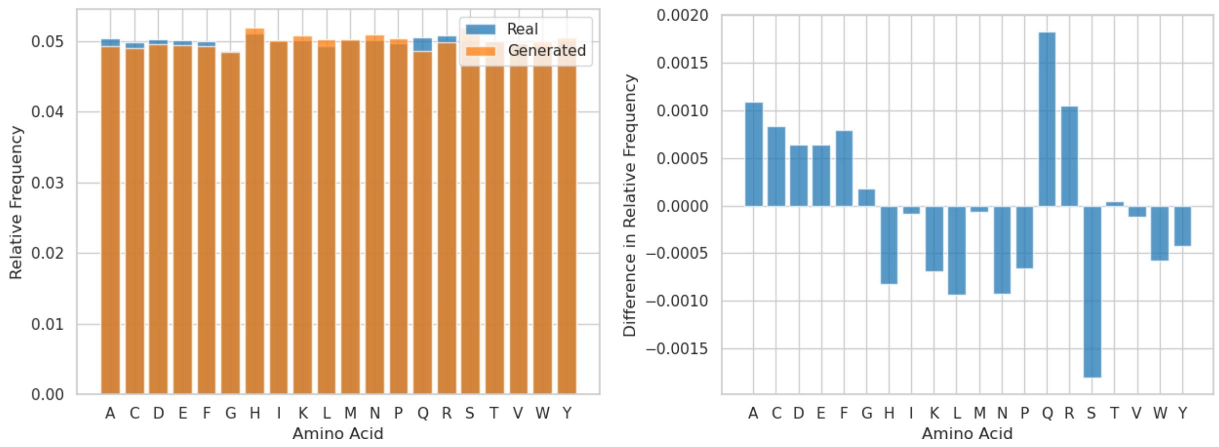


Figure S7: Amino acid frequency distribution comparison between two independent groups of 5000 uniformly randomly constructed sequences with a maximum length of 32. The distributions are flat, excluding a small amount of sampling noise. Additionally, the deviation between the two is extremely small, with the largest difference value being several orders of magnitude smaller than the largest value present in Figures 3 or S6.

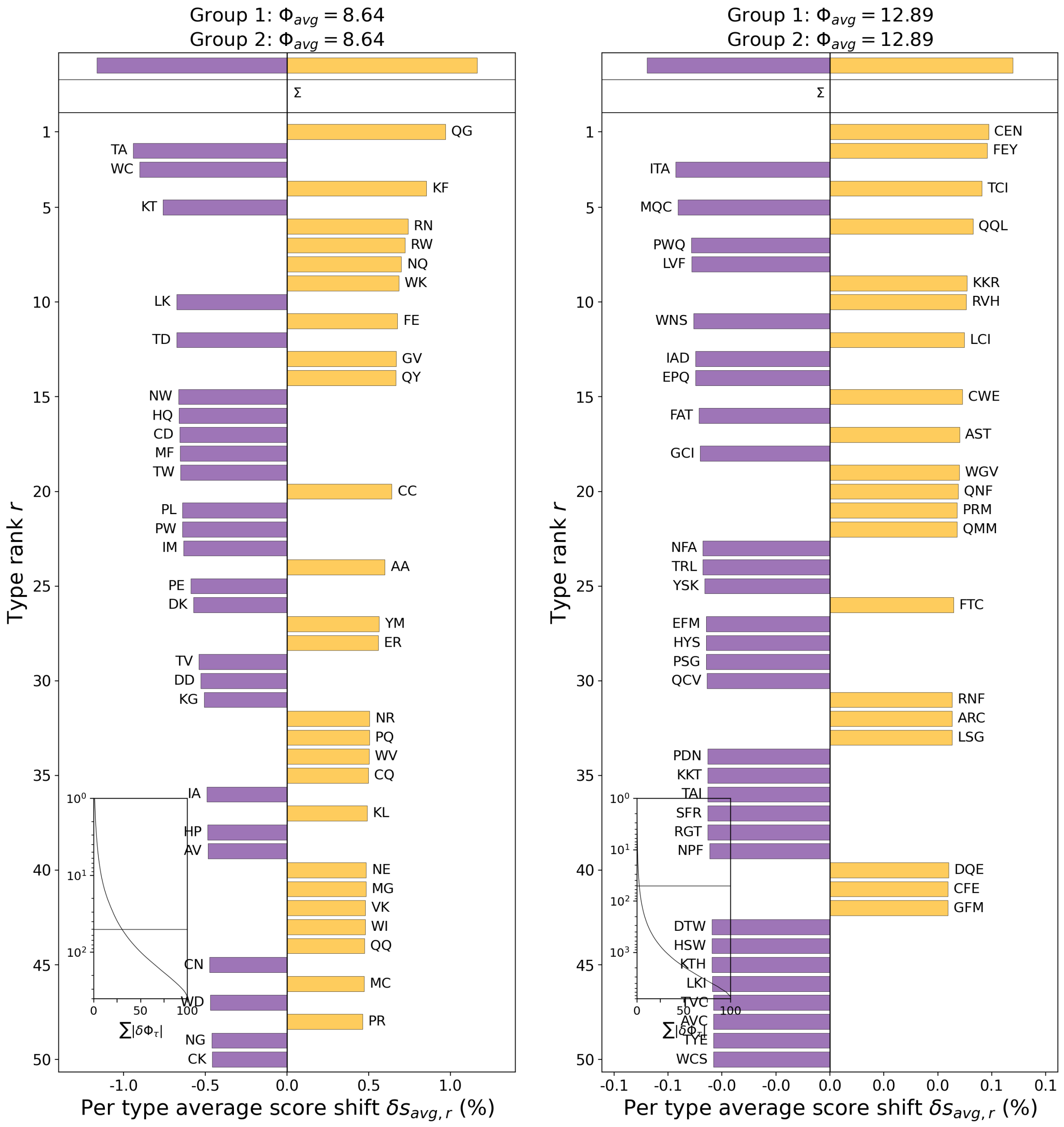


Figure S8: Word shift plots comparing two independent groups of 5000 uniformly randomly constructed sequences with a maximum length of 32. Similar to the character level analysis shown in Figure S7, these word shifts are extremely flat. However, since the number of distinct elements grows exponentially with the sub-sequence length, sampling error may have a larger impact here. The maximum entropy for length 2 sub-sequences constructed from the 20 common amino acids is ∼8.64, which is reliably obtained by a sample of this size. The maximum entropy for length 3 sub-sequences is ∼12.97, but is not reached due to sampling error. Approximately 1 to 5 length 3 sub-sequences are unobserved in a sample of this size. There are 400 unique length 2 and 8000 unique length 3 sub-sequences, thus a uniform distribution over those sets has an element-wise probability of 0.0025 and 0.000125 respectively.

### Global Sequence Alignment Scores

To investigate the similarity of two bags of sequences we applied the Gotoh global alignment algorithm.[78] We use the implementation provided by Biopython's `PairwiseAligner` object,[79] configured with the `BLOSUM62` substitution matrix, an open gap score of -10, and an extend gap score of -1.

Interpreting global alignment scores can be difficult, so we performed a Monte Carlo experiment to uncover information about the distribution of scores in particular circumstances. Specifically, we construct two bags of random sequences, called $S_1$ and $S_2$, containing $N$ and $N/2$ sequences respectively. These sequences have a uniformly random length selected from 1 to 32, and the elements of each sequence are uniformly randomly selected from the 20 common amino acids. Next, we construct a new bag, $S_3$, by combining $S_2$ with a duplicate. Thus, $S_1$ and $S_3$ both contain $N$ sequences, where all sequences in $S_1$ are likely to be unique and $S_3$ contains two copies of every unique sequence in $S_2$. Next, we construct $S_4$ by randomly mixing the sequences of $S_1$ and $S_3$ using a control parameter $m \in [0, 1]$. This mixing is implemented by iterating pairwise over the sequences of $S_1$ and $S_3$, then iterating pairwise over the FASTA characters of those sequences. The characters from $S_1$ are selected with probability $1 - m$ and the characters from $S_3$ are selected with probability $m$. From this construction, the mixing parameter directly controls the diversity of $S_4$, providing a stochastic interpolation between relatively high and low diversity bags of sequences. Finally, pairwise scoring is computed between $S_4$ and itself using the system described above.

The full experiment then involved sampling the mixture parameter at 50 evenly spaced points that span the interval $[0, 1]$ and executing 30 replicates of the scoring procedure at each. All replicates use $N = 1000$. Figure S9 summarizes this experiment, where the orange line indicates the mean match score across the replicates and the shaded area covers plus or minus one standard deviation. Percentile information is shown by the blue lines, and extreme values are shown with grey markers. This indicates a roughly exponential scaling in the expected match score as the mixing parameter varies from 0 to 1, ranging from ∼2.5 to ∼20. The standard deviation indicates the existence of heteroscedasticity, where the tails of the score distribution spread out as the mixture parameter increases in value.

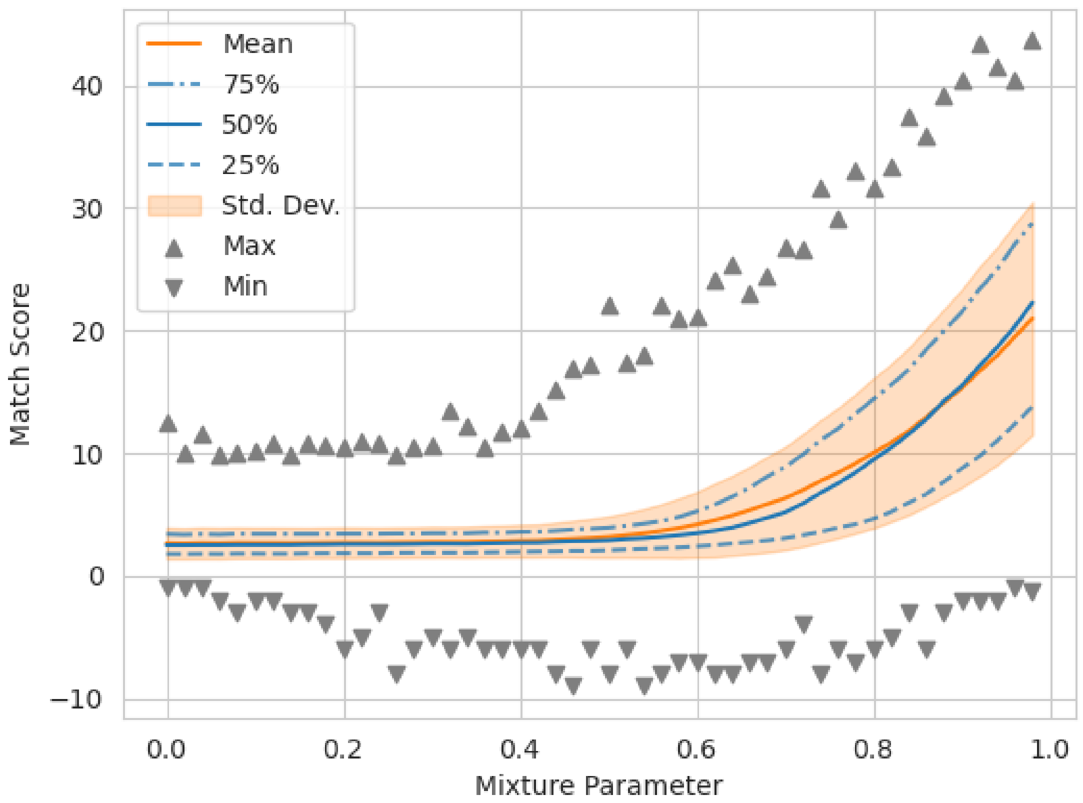


Figure S9: Distributions of global match scores between a bag of FASTA sequences and itself. The dark line indicates the mean match score and the shaded area indicates plus or minus one standard deviation. The horizontal axis corresponds with a mixture parameter that controls the level of diversity in the bag. For low values the bag of sequences is composed entirely of unique sequences, resulting in low match scores on average. As the value of the mixture parameter increases the level of diversity in the bag decreases. When the mixture parameter reaches a value of 1.0 the bag contains an exact duplicate for every sequence, resulting in match scores in the 30s.
